# Stable integrant-specific differences in bimodal HIV-1 expression patterns revealed by high-throughput analysis

**DOI:** 10.1101/666941

**Authors:** David F. Read, Edmond Atindaana, Kalyani Pyaram, Feng Yang, Sarah Emery, Anna Cheong, Katherine R. Nakama, Erin T. Larragoite, Emilie Battivelli, Eric Verdin, Vicente Planelles, Cheong-Hee Chang, Alice Telesnitsky, Jeffrey M. Kidd

**Affiliations:** Departments of Human Genetics, University of Michigan Medical School, Accra, Ghana; Departments of Microbiology and Immunology, University of Michigan Medical School; Accra, Ghana; Departments of West African Centre for Cell Biology of Infectious Pathogens (WACCBIP), Department of Biochemistry, Cell & Molecular Biology, University of Ghana, Accra, Ghana; Department of Pathology, University of Utah, Salt Lake City, UT; Department of Medicine, University of California San Francisco, San Francisco, CA; Buck Institute for Research on Aging, Novato, CA

## Abstract

HIV-1 gene expression is regulated by host and viral factors that interact with viral motifs and is influenced by proviral integration sites. Here, expression variation among integrants was followed for hundreds of individual proviral clones within polyclonal populations throughout successive rounds of virus and cultured cell replication. Initial findings in immortalized cells were validated using CD4+ cells from donor blood. Tracking clonal behavior by proviral “zip codes” indicated that mutational inactivation during reverse transcription was rare, while clonal expansion and proviral expression states varied widely. By sorting for provirus expression using a GFP reporter in the *nef* open reading frame, distinct clone-specific variation in on/off proportions were observed that spanned three orders of magnitude. Tracking GFP phenotypes over time revealed that as cells divided, their progeny alternated between HIV transcriptional activity and non-activity. Despite these phenotypic oscillations, the overall GFP+ population within each clone was remarkably stable, with clones maintaining clone-specific equilibrium mixtures of GFP+ and GFP-cells. Integration sites were analyzed for correlations between genomic features and the epigenetic phenomena described here. Integrants inserted in genes’ sense orientation were more frequently found to be GFP negative than those in the antisense orientation, and clones with high GFP+ proportions were more distal to repressive H3K9me3 peaks than low GFP+ clones. Clones with low frequencies of GFP positivity appeared to expand more rapidly than clones for which most cells were GFP+, even though the tested proviruses were Vpr-. Thus, much of the increase in the GFP-population in these polyclonal pools over time reflected differential clonal expansion. Together, these results underscore the temporal and quantitative variability in HIV-1 gene expression among proviral clones that are conferred in the absence of metabolic or cell-type dependent variability, and shed light on cell-intrinsic layers of regulation that affect HIV-1 population dynamics.

**Summary:** Very few HIV-1 infected cells persist in patients for more than a couple days, but those that do pose life-long health risks. Strategies designed to eliminate these cells have been based on assumptions about what viral properties allow infected cell survival. However, such approaches for HIV-1 eradication have not yet shown therapeutic promise, possibly because much of the research underlying assumptions about virus persistence has been focused on a limited number of infected cell types, the averaged behavior of cells in diverse populations, or snapshot views. Here, we developed a high-throughput approach to study hundreds of distinct HIV-1 infected cells and their progeny over time in an unbiased way. This revealed that each virus established its own pattern of gene expression that, upon infected cell division, was stably transmitted to all progeny cells. Expression patterns consisted of alternating waves of activity and inactivity, with the extent of activity differing among infected cell families over a 1000-fold range. The dynamics and variability among infected cells and within complex populations that the work here revealed has not previously been evident, and may help establish more accurate correlates of persistent HIV-1 infection.

## Introduction

Early in the HIV-1 replication cycle, a DNA intermediate integrates into the host cell’s genome. HIV-1 replication ordinarily progresses into its late phases, with host and viral factors leading to gene expression, virion production, and cell death and/or virus spread. However, some proviruses can remain dormant upon integration. In patients, the resulting latently infected cells persist throughout antiretroviral treatment, and their sporadic reactivation can lead to virus rebound after antiretroviral cessation.

This source of persistent virus is called the latent reservoir, and is believed to consist largely of transcriptionally silent proviruses integrated into resting memory T cells [1] [2] [3]. Experimentally, infectious virus can be produced from such patients’ T lymphocytes when they are activated or treated with certain chromatin remodeling drugs *ex vivo*. These observations inspired “shock and kill” HIV cure strategies, which involve pharmacologically inducing provirus expression to promote the recognition and clearance of latently infected cells [4] [5]. However, while drug treatments that reactivate silenced proviruses can activate HIV-1 gene expression in cell culture models of latency, such treatments have thus far failed to fulfill their promise in the clinic, suggesting much remains to be learned about the establishment and maintenance of the latent reservoir [6] [7] [8].

HIV-1 gene expression requires sequence motifs within proviral sequences that specify nucleosome positioning and allow HIV-1 to respond to host factor differences among infected cell types [9] [10] [11]. HIV-1 has a marked preference for integration in transcriptionally active genome regions [12, 13]. Specific integration sites also influence HIV gene expression [14] [15] [16] [17]. Certain host chromatin binding factors as well as nuclear architecture further bias the distribution of integration sites [18, 19]. It has been postulated that integration sites may affect the odds of a provirus establishing long-lived latency [20]. Differences in HIV-1 expression due to integration site features likely influence the extent to which cells survive and proliferate after HIV-1 integration, and in turn contribute to the expression profile of persistent HIV-1 [21].

Recent work with patient samples has demonstrated that for at least some suppressed patients, residual provirus-containing cells are polyclonal yet dominated by a limited number of clonal subsets [22], and similar observations of clonal expansion have been made during HIV-1 infection of humanized mice [23]. Thus, the integration sites represented in persistent proviruses likely differ from the spectrum initially generated [21].

Recent evidence indicates that latent proviruses differ in the extents to which they can be reactivated, and that a large majority of cells harboring latent proviruses may be refractory to our current arsenal of reactivation agents [24, 25]. Work using dual color reporter viruses in primary cells has shown that proviruses differ in their reactivation potential depending on their sites of integration, with chromatin context as maintained within the confines of the nucleus being a significant contributing factor [25]. Additional work monitoring HIV-1 expression in individual cells has questioned the earlier view that complete proviral silencing is necessary for infected cell persistence during antiretroviral therapy [26, 27].

The majority of proviruses detectable in suppressed patients are replication defective [26, 28]. Although such proviruses are incapable of rekindling infection, emerging evidence suggests they can be expressed and may contribute to pathogenesis [26, 29].

In this study, we developed a high throughput approach to monitor cellular and viral progeny of individual integration events within complex populations, and used it to address the frequency of defective provirus formation and the extent to which provirus integration sites affect provirus expression levels. Initial work was performed using transformed cell lines, where selective pressures and variation of intracellular factors should be lower than in primary cells, with additional experiments performed in CD4+ lymphocytes from donor blood. Examining the extent of expression variation within and among cellular progeny of large panels of individual HIV-1 integration events indicated that in all these cell types, epigenetic differences among proviral clones led to the establishment of distinct heritable patterns of HIV-1 gene expression.

## Results

### Using randomized vectors to produce zip coded proviruses

We developed a system to uniquely identify individual HIV-1 proviral lineages within polyclonal integrant populations, track proviral gene expression, and monitor replication properties of individual cell clones and their viral progeny. To achieve this, HIV-1 based vectors were established that each contained a unique 20-base randomized sequence tag. Once integrated, these were referred to as “zip coded” proviruses because the randomized tags reported the proviruses’ locations in the human genome.

Zip coded proviruses were templated by pNL4-3 GPP, a NL4-3 strain derivative that encodes Gag, Pol, Tat and Rev plus a puromycin-resistance reporter expressed from a secondary, internal promoter (SV40; Figure 1A, upper construct). pNL4-3 GPP lacks *env* and all accessory genes including *nef*, which was disrupted by the sequence tags [30]. Tags were inserted into the upstream edge of U3, downstream of sequences required for integrase recognition and upstream of the site of nuc0 nucleosome binding [11]. Vector RNAs were transcribed from uncloned DNA template libraries containing randomized tags, which were generated by *in vitro* assembly and without amplification by plasmid replication. This experimental approach resulted in the introduction of a unique randomized 20-mer into each encapsidated viral RNA. Because the process of reverse transcription duplicates U3, each progeny provirus contained the same randomized tag in both LTRs, and each provirus’s tags differed from those in every other integrant.

**Figure 1.**
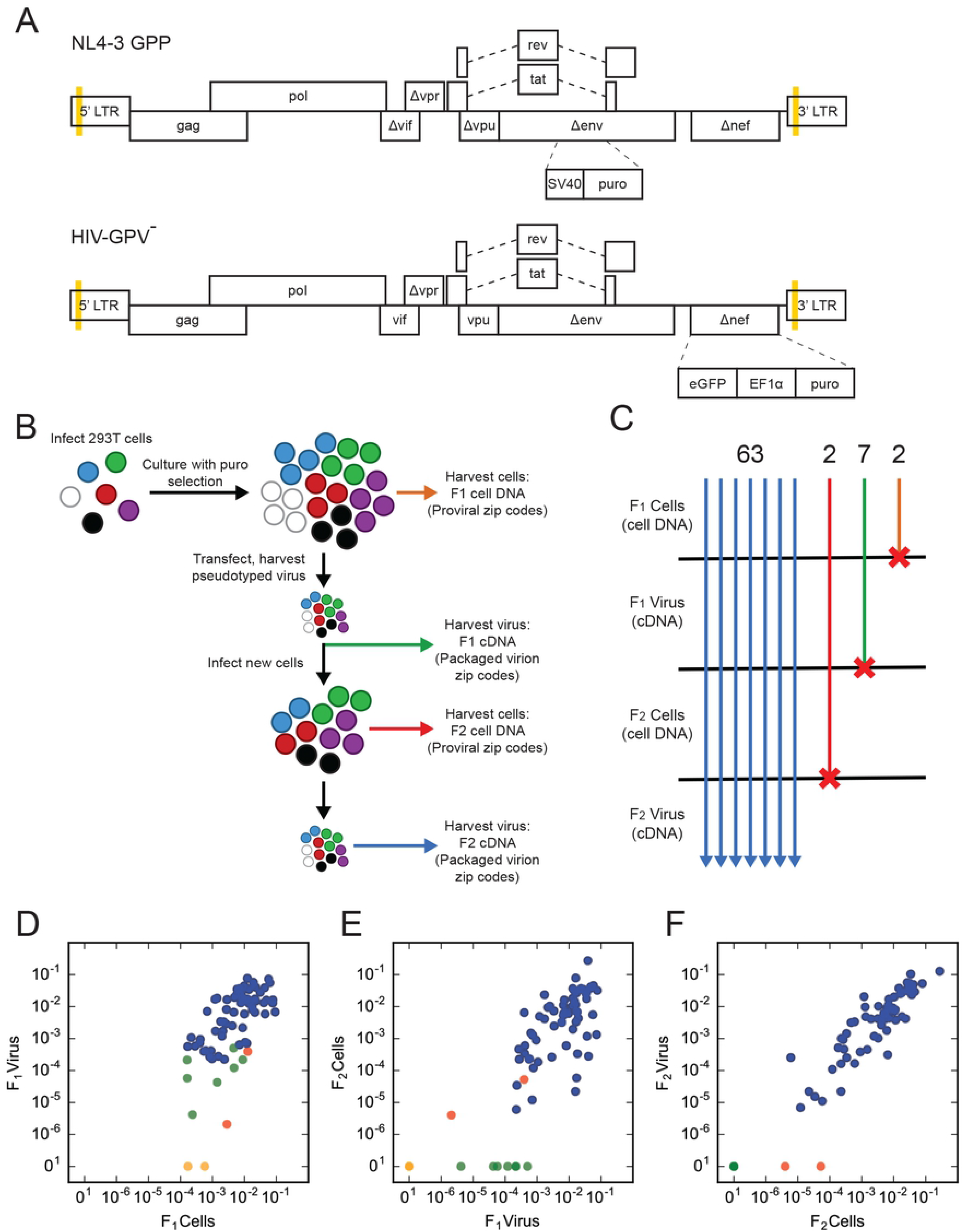
Monitoring proviral replication competence across generations. A: Schematic illustrations of the vectors used in this paper. Gold bars represent the sites of randomized sequence insertions. Features and construction are described in Materials and Methods. B: A schematic of the experimental flow of the replication competence experiment, depicting the analysis of genomic DNA and viral cDNA harvested from the F_1_ and F_2_ generations. C: Summary of the number of independent zip codes detectable at different stages of the experiment. A total of 63 zip codes were detected in all four pools. The remainder were only detected in the indicated stages. D-F: scatter plots of zip code read proportions across indicated stages of the experiment, as outlined in panel B. Each clone is represented by a single point, colored to reflect that clone’s persistence based upon the progression pattern depicted in panel C. The Spearman correlation for each comparison is given.

To validate this approach, adherent 293T cells were transduced at a very low (<0.00005) multiplicity of infection with VSV-G pseudotyped particles containing a random sequence-containing vector library. Ten individual puromycin-resistant colonies were expanded, proviral DNA was amplified, and PCR products encompassing the randomized region were sequenced. The results showed that one clone lacked an insert while each of the other nine contained a single unique randomized 20-mer, no two of which were the same (Supplementary Table 1).

### Nearly 90% of first-round proviruses supported a second round of replication

An initial pooled-clone experiment was then performed, which established analysis procedures using a proviral pool of known complexity. This pilot study also addressed the frequency of defective provirus formation during a single replication round (Figure 1B). The pool was generated by infecting 293T cells with virions containing zip coded NL4-3 GPP RNAs, selecting a total of 71 well-separated puromycin-resistant colonies, trypsinizing them, and pooling these cells to generate the so-called F_1_ cell pool. After expansion, a portion of the F_1_ cell pool was transfected with a VSV-G expression plasmid to generate pseudotyped virions (“F_1_ virus”) that were used to infect fresh 293T cells. These “second generation” F_2_ cells were placed under selection pressure and the resulting colonies were pooled to generate the F_2_ cell pool. Because the number of colonies pooled to generate the F_2_ cell pool--roughly 1000-- was significantly greater than the F_1_ pool’s zip code complexity, any infectious zip code present in the F_1_ pool was predicted to generate multiple F_2_ integrants.

The ability of each provirus generated during the first round of replication to successfully complete a second round was addressed by examining each zip code’s prevalence throughout experimental stages. RNA from F_1_ and F_2_ virions was harvested and used to template cDNA libraries, and genomic DNA extracted from F_1_ and F_2_ cell pools was used to make provirus zip code libraries. High throughput data from all four libraries was analyzed to compare the zip code content of each pool (Figure 1B).

DNA from F_1_ cells was found to contain 74 unique zip codes, which accounted for 99.87% of total sequencing reads (Supplementary Figure 1). Based on the low multiplicity of infection used here and on subsequently-determined differences among zip codes in expression properties, the discrepancy between this value and the 71 colonies visualized on tissue culture plates was likely due to miscounting double colonies as single expanded clones, although the possibility of dual infection cannot be ruled out. F_1_ cell, F_1_ virus, and F_2_ cell libraries were then compared to determine how many proviruses remained infectious after their initial round of replication (Figure 1C). Because 65 out of the 74 zip codes found in F_1_ cell DNA were also observed in the F_2_ cell library, these 65 (88% of F_1_ cell zip codes) unambiguously represented proviruses capable of completing a second round of replication.

The remaining 9 zip codes were candidate non-infectious proviruses. If a first-round provirus were defective in ways that allowed virion assembly but not replication, such virus’ zip code might be detectable in F_1_ virus but not in F_2_ cells. Seven zip codes were candidates for this class of defective proviruses (green lines in Figure 1C).

The remaining two clones were initially enigmatic. The total number of colonies pooled to generate the F2 library suggested it contained roughly twenty re-transduced copies of each F1 zip code. Based on how frequently replication competence was maintained after the first round of replication, it was expected that any fully infectious F_1_ provirus would display a roughly 90% second-round success rate. Thus, the likelihood that all ∼20 sibling F_2_ progeny of any infectious F_1_ provirus would be defective seemed exceptionally low. Incongruously however, among the 65 replication-competent zip codes detected in the F_2_ cell library, two were not observed among sequencing reads from the F_2_ virus RNA library.

### Integrant clone expansion and provirus expression levels varied widely among zip coded 293T cell clones

To address whether the absence of two F_2_ cell zip codes from the F_2_ virus library might reflect a population bottleneck, the number of sequencing reads associated with each zip code was compared within and across libraries. Unexpectedly, reads per zip code were observed to vary over three orders of magnitude within the F_1_ cell library (Figure 1 D). Although variations in provirus-containing cells’ expansion rates have been reported previously [31], the wide range in cell clone sizes observed here had not been anticipated.

Clone-specific differences in the amount of virus released per cell were also observed (Figure 1D, y axis). When normalized to the number of F_1_ cells harboring a given zip code, clone-associated differences in virion release per cell spanned two orders of magnitude or more. Because of this, zip code abundance in the F_1_ cell library was only moderately correlated with abundance in the F_1_ virus RNA library (Figure 1D) (Spearman ρ = .639). In contrast, the correlation between cell count and virion production was strong in the F_2_ generation (Spearman ρ = .890) where each zip code was polyclonal (Figure 1 F), suggesting that virus-per-cell ratios were fairly consistent when averaged across many cell clones.

Looking specifically at sequencing read data for the two F_2_ cell zip codes that were missing from F_2_ virus libraries revealed that these lineages were scarce in both the F_1_ virus and in F_2_ cells (red points, Figure 1 E). Similarly, read frequency trends for the seven F_1_ zip codes not observed in F_2_ cells (green points, Figure 1 D) suggested that population bottlenecks, and not loss of infectivity, may account for some of these candidate non-infectious zip codes’ absence from F_2_ cells.

### Clonal expansion in Jurkat cells

Larger zip coded integrant populations were then established using Jurkat cells. The vector in these experiments (HIV GPV**^-^**) expressed all HIV-1 genes except *env*, *nef*, and *vpr*, contained GFP in the *nef* open reading frame, and expressed a puromycin resistance marker from a secondary, internal promoter (Figure 1A, lower construct). Selective concentrations of puromycin were applied 24 hours post-infection and removed four days later. Cells were subsequently maintained without drug. Like dual color vectors where one marker is LTR-driven and the other is driven by a secondary promoter, this approach used a marker expressed from a secondary promoter to identify productively infected cells, independent of the LTR’s transcription state [32]. However, because puromycin is labile and drug selection was applied briefly, the analysis here may have included clones for which the entire provirus, including the internal promoter, was silenced after an initial period of activity. Cell pools infected at differing multiplicities were analyzed by high throughput sequencing, and a Jurkat pool comprised of roughly 1,000 zip coded clones was used in subsequent studies.

The sequencing of zip codes amplified from duplicate aliquots of this pool’s cells revealed the presence of many zip codes shared in both replicates. However, lower abundance zip codes were sampled unevenly. To better address the pool’s complexity and differential clone expansion, ten technical replicates made from the pool’s genomic DNA were combined to provide evidence for 706 zip codes, which together accounted for 97.8% of total reads and displayed clonal abundances spanning over two orders of magnitude (Figure 2, Supplementary Figure 2).

**Figure 2.**
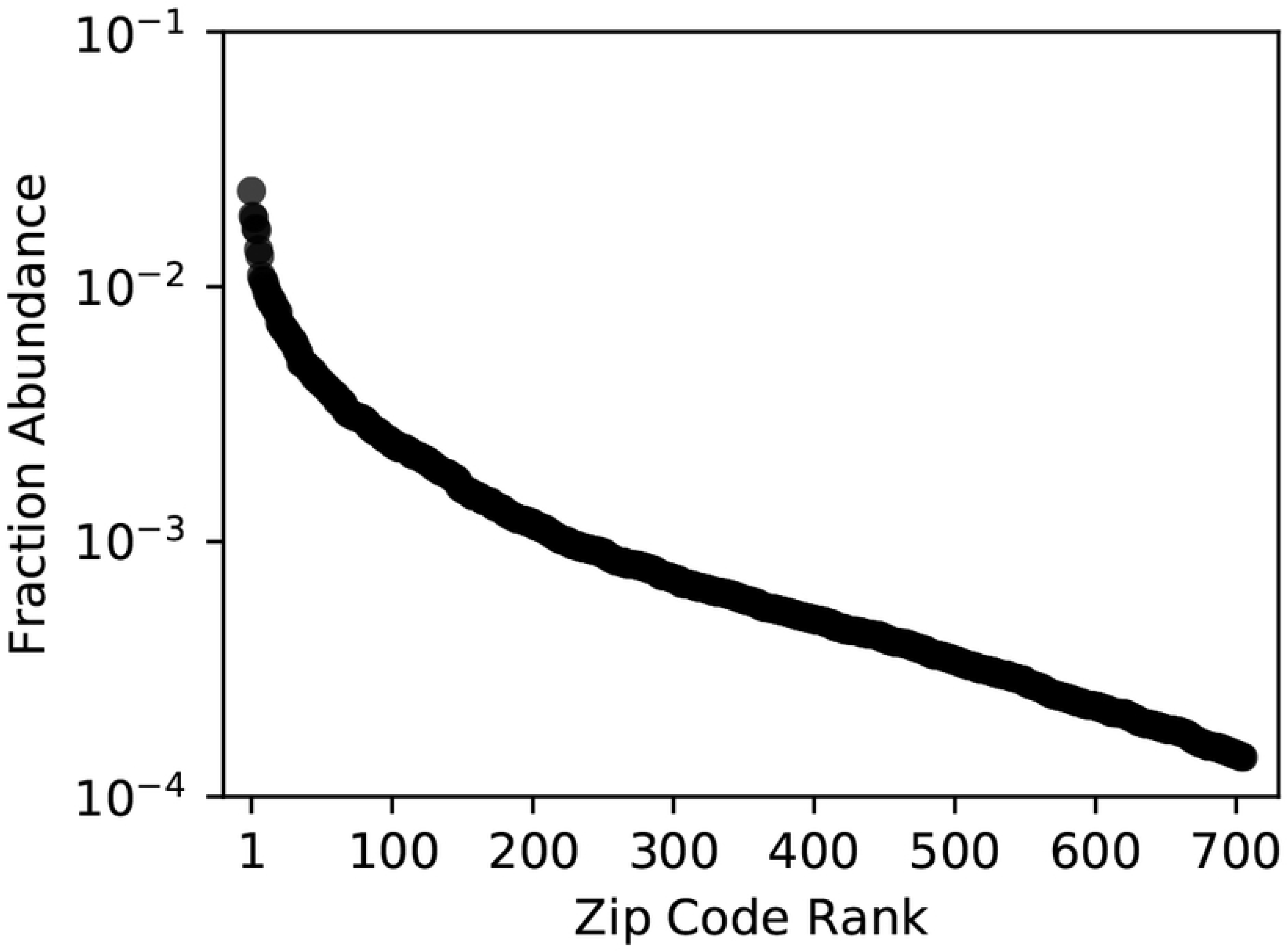
Zip code fractional abundance. Each of 706 zip code families identified in the Jurkat pool is depicted by a single point. The clones are arrayed left to right from the most abundant to the least abundant, with the fractional abundance of total reads assigned to that zip code on the Y axis.

### Significant clone-by-clone differences in HIV-1 expression in both Jurkat and primary cells

Detecting GFP by flow cytometry allowed binary (on/off) monitoring of LTR expression in individual cells. Portions of the total Jurkat pool, designated Pools 1 and 2, were independently sorted into GFP positive “GFP+” and negative “GFP-” sub-pools (Figure 3A). The zip code content of each sub-pool was assessed by high throughput sequencing, to simultaneous quantify expression characteristics of hundreds of clonal lines (Figure 3). As a control, we also sorted cells based on their p24 content, using an anti-p24 antibody, and observed a strong correlation between GFP expression and p24 content. (Supplementary Figure 3).

**Figure 3.**
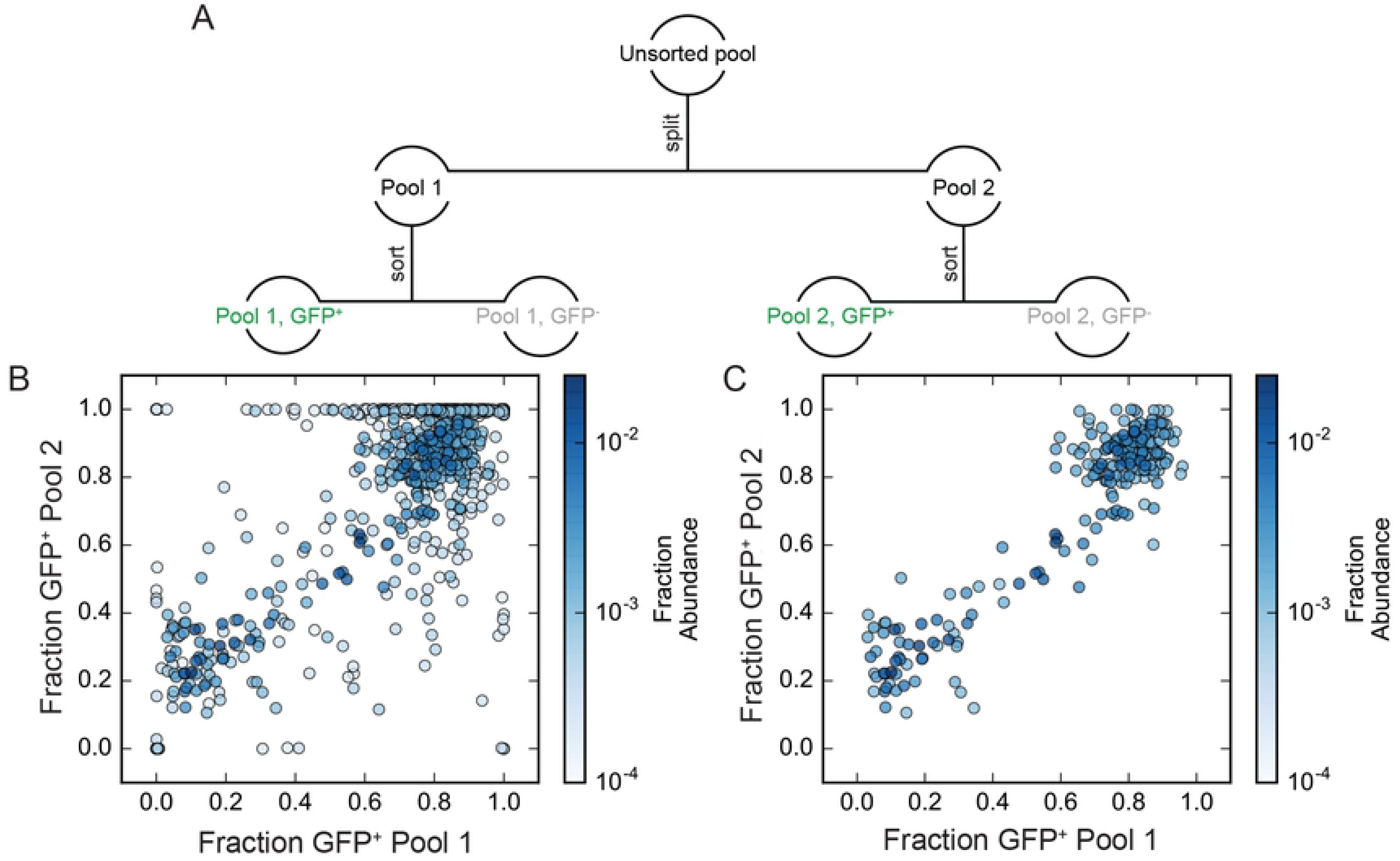
GFP+ proportions for independent clonal lines within a complex population. A: Schematic description of the cell pool splitting and sorting procedures performed. GFP+ proportions were determined as described in the text. B: comparison of fraction GFP+ determined for each zip code in pool 1 and pool 2. Each point represents a single zip coded cell clone. Individual clones are colored based on their fractional abundance in the original unsorted pool as indicated by the color bar at the panel’s right. C, as in B, but with data for the less abundant clones removed to show only the 225 zip codes with fractional abundance > 0.001.

An expression value termed the “GFP+ proportion” was determined for each zip coded clone. GFP+ proportions were calculated by dividing the read frequency of each zip code within GFP+ sorted cells by the zip code’s summed abundance in both GFP+ and GFP-sorted cells. To accurately represent zip codes’ prevalence in the total pool, GFP+ and GFP-clone abundance values were weighted to reflect the proportions of total Pool 1 and 2 cells contained in GFP+ and GFP- sub-pools. A sample calculation is provided in Materials and Methods. Consistent with clonal variation in virus release per cell observed in the pilot experiment above, GFP+ proportions differed significantly among Jurkat cell clones, with individual clones’ GFP+ proportions ranging from >99% to <1%.

To better understand if the broad range of GFP+ proportions observed among clones reflected clone-specific properties or were a result of sampling, we compared data for duplicate experimental samples, with the GFP+ proportions calculated for each zip code in Pool 1 compared to GFP+ proportions independently determined for Pool 2. As shown in Figure 3B, when GFP+ proportion data were plotted against each other, most clones displayed similar values, suggesting that each clone possessed a distinct GFP+ proportion that was not defined by sampling (Spearman ρ = 0.474 for the 688 zip codes detected in each pool). GFP+ proportions were particularly well correlated for the most abundant zip codes (Figure 3C, Spearman ρ = 0.716 for the 225 zip codes with fractional abundance > 0.001 in the parental pool), suggesting that at least 200 clones were sufficiently abundant in the total population to be reproducibly well sampled in repeated sub-pools.

Both experiments above were performed with cell lines, where within-experiment differences in environment and *trans-*acting factors should be minimized [33]. In an initial test of whether primary cells also displayed integrant-specific differences in our system, CD4+ cells were isolated from donor blood, stimulated, and transduced with VSV-G pseudotyped zip coded GPV**^-^** (Figure 4). A low multiplicity of infection was selected to aid analysis based on the limited cell divisions that occur after a single round of primary cell stimulation.

**Figure 4.**
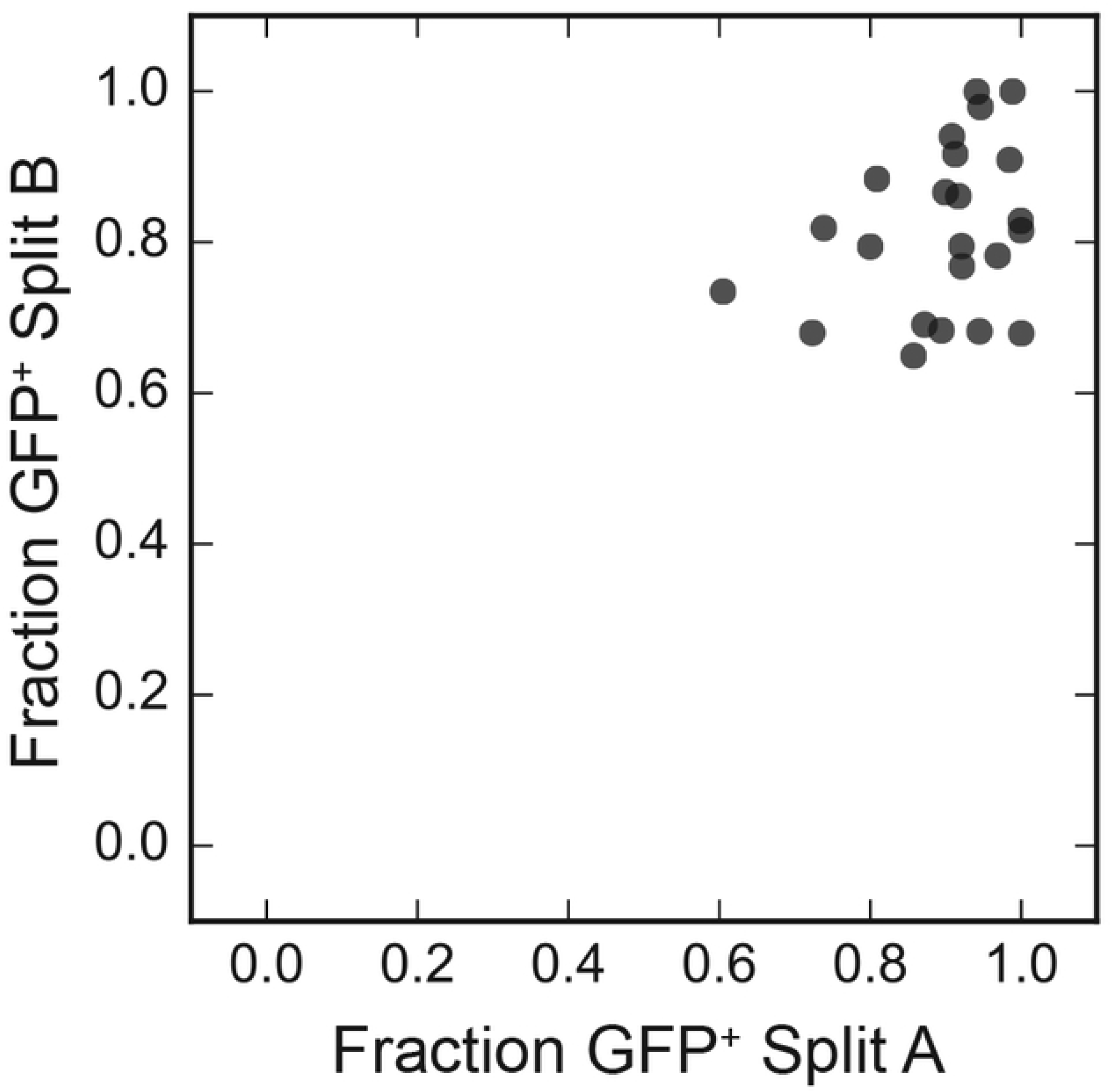
Reproducibility of GFP+ proportions for individual primary cell clones. The GFP+ proportions for 24 zip codes selected as described in the text and detected in two sub-pools of a primary cell infection were plotted against one another.

Six days post infection, the cells were divided into 2 sub-pools and then sorted into GFP- and GFP+ cell fractions. Genomic DNA was isolated from both sub-pools’ cell fractions, and proviral zip codes were amplified and sequenced. These experiments were performed in the absence of puromycin selection. Thus, the GFP-fraction included both uninfected cells and cells containing silent proviruses, and both the first and the second sub-pools displayed very low levels of GFP+ cells (0.62% and 0.52% respectively). As a result, precise GFP+ proportions could not be determined because the total infected cell frequency was not measured, and instead an assumed frequency of 10% GFP-cells was used in Figure 4 calculations, based on parallel control experiments performed at a higher MOI and with puromycin selection. Note that this 10% value is on the upper edge of previously reported primary cell values and may reflect donor-dependent variation or survival of some non-transduced cells [34]. However, although absolute values would change if true GFP-value were lower than assumed, correlation trends and their interpretation would not be affected.

Consistent with the results observed with 293T and Jurkat cells, sequencing read data suggested significant differences among primary cell integrants in clonal expansion rates, as their progeny cells’ abundance levels spread over a wide range (Supplementary Figure 4). Due to the relatively short duration of primary cell propagation, these abundance differences severely curtailed the number of clones that were sampled sufficiently to meet inclusion criteria (that the clone was detectable with fractional abundance > 0.001 in each sub-pool). Nonetheless, when the GFP+ proportions for these primary cell zip codes were calculated for each independently analyzed sub-pool and values for the two replicate sorts were plotted against one another (Figure 4), the trends significantly supported the likelihood that provirus-containing progeny of primary cells differed by clone in their levels of HIV-1 gene expression (Spearman ρ = 0.26).

### Clones’ GFP+ proportions are a stable, heritable phenotype

Longitudinal studies were performed with the zip coded Jurkat pool to monitor GFP+ proportions throughout cell generations. After sampling for sequencing library preparation, aliquots of the two GFP+ and two GFP- sub-pools analyzed in Figure 3A were passaged separately for an additional 8 to 9 days, at which time point each of these four pools was again sorted by FACS (Figure 5). The results showed that the cellular descendants of Pool 1 and Pool 2 GFP+ sub-pools did not all remain GFP+, nor did the descendants of the GFP- sub-pools remain all GFP-. Instead, some cells from each sub-pool had switched expression phenotypes during passaging. This suggested that the HIV-1 expression pattern in any individual cell was not stably inherited by all of its progeny, but that instead expression “flickered” (alternated between LTR expression and silencing) during cell propagation.

**Figure 5.**
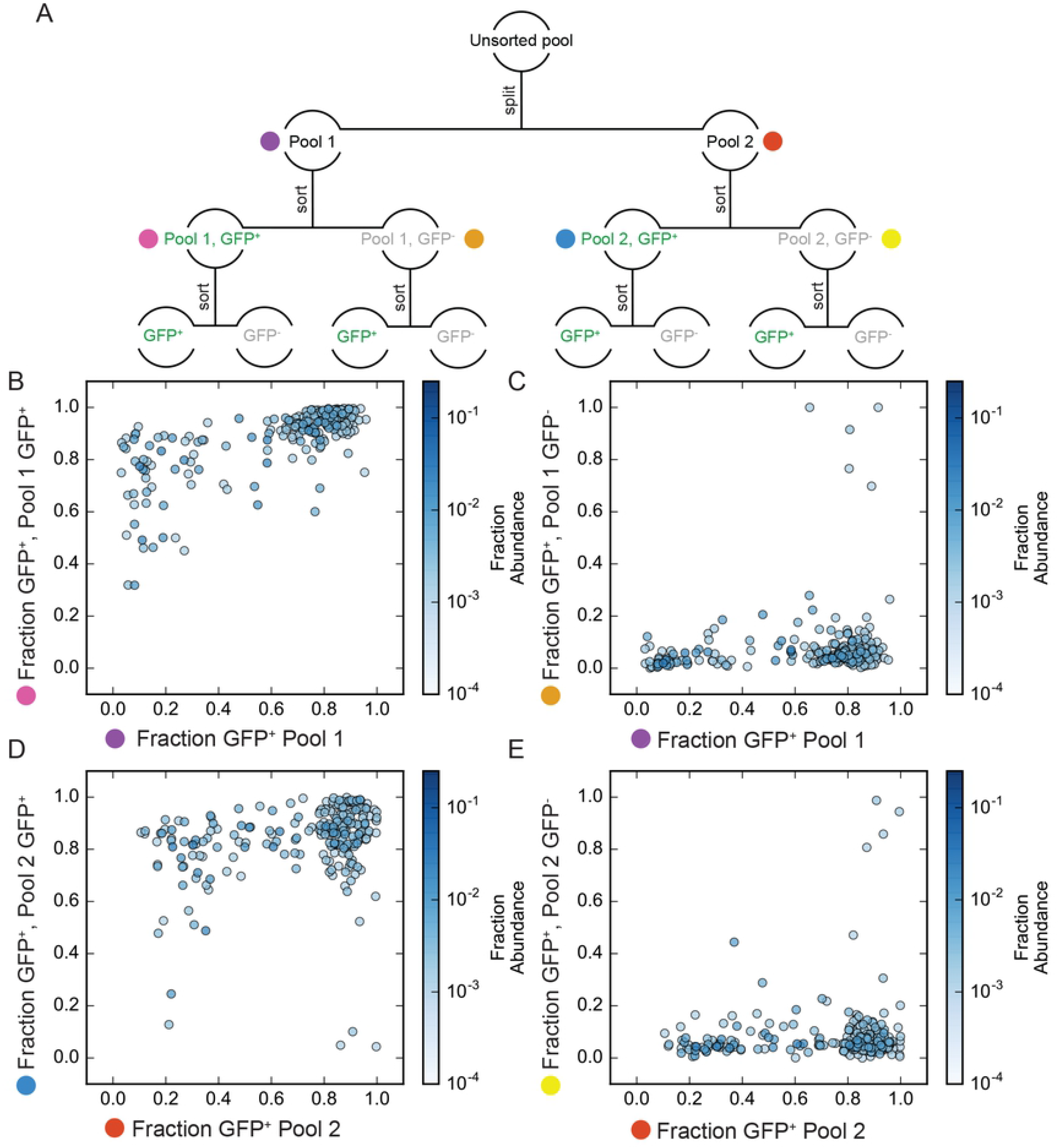
GFP+ proportions of passaged and re-sorted GFP+ and GFP-cell pools. A: Depiction of the cells’ passaging and sorting scheme, with the initial sorted pools characterized in Figure 3 at the top, followed by the re-sorted sub-pools analyzed here. The colored dots next to each pool correspond to the comparisons plotted in panels B-E. B: Analysis of zip codes that sorted GFP+ in pool 1. C: Analysis of zip codes that sorted GFP-in pool 1. D: Analysis of zip codes that sorted GFP+ in pool 2. E: Analysis of zip codes that sorted GFP-in pool 2

Integrant specific, intrinsic rates of expression that are maintained across cell generations have previously been reported for basal expression from the HIV-1 promoter [13]. To test whether or not the expression patterns studied here also were stable over time, the GFP+ proportions determined for the GFP+ or GFP-pools in the second sort were combined after weighting to reconstitute the parental pool (Pool 1 or 2 in Figure 3A) proportions. As described in Materials and Methods for the derivation of GFP+ proportion values, these adjustments allowed frequencies determined within individual sub-pools to be combined in a way that accounted for the composition of the original unfractionated population. This was especially important here, where the first-sort GFP+ sub-pool had been heavily enriched for cells from clones with high GFP+ proportions, while the first-sort GFP- sub-pools were enriched for clones with low GFP+ proportions. Consistent with the stable inheritance of clone-specific intrinsic expression patterns, the data indicated that the weighted GFP+ proportions for each integrant following the second sort showed a strong correlation with its original GFP+ proportion (Spearman ρ = 0.938 for Pool 1 and 0.805 for Pool 2; Figure 6).

**Figure 6.**
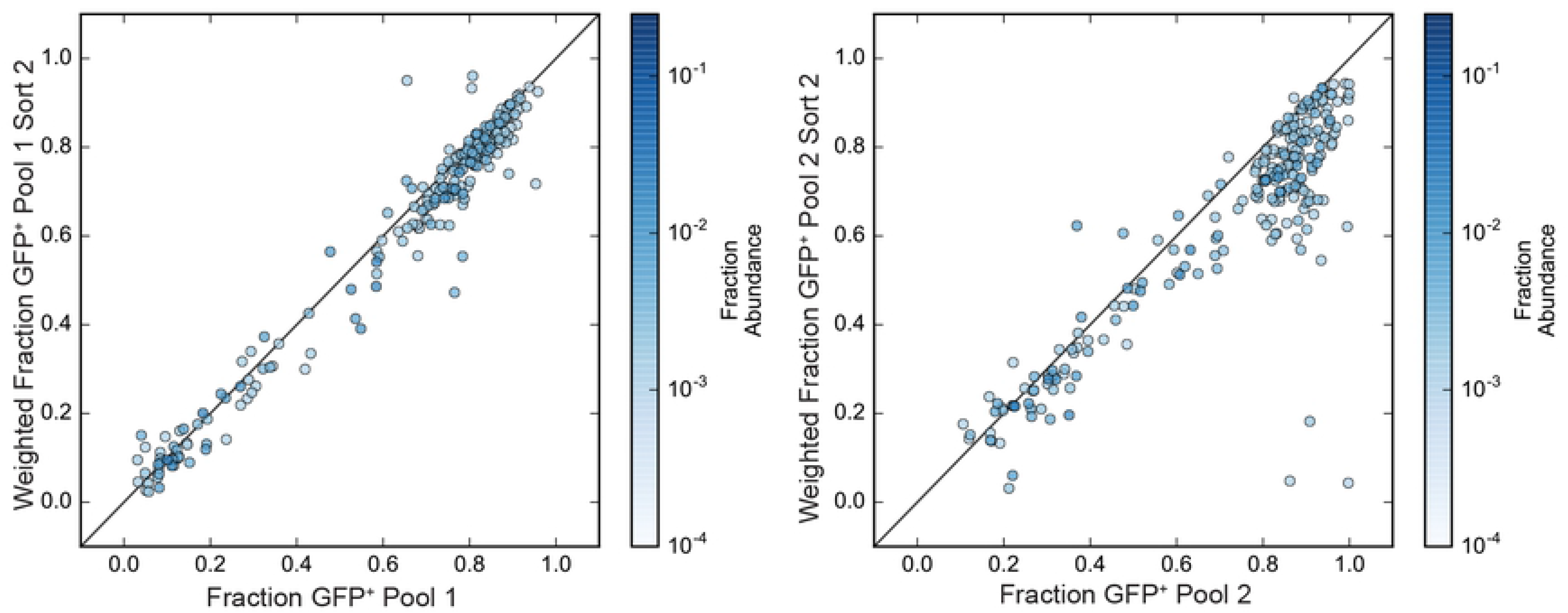
Stability of GFP+ proportions over time. GFP+ proportions determined in the first sort (Figure 3 data) plotted against reconstructed GFP+ proportions for each zip code derived from data in the second sort (Figure 5). Second sort GFP+ proportions were reconstructed by weighting the GFP+ and GFP- sub-pool values determined in Figure 5 and combining these to reconstitute GFP+ proportions for the total pool after extended passage, and these second sort reconstituted proportions were plotted against experimental values from the first sort.

### Integration site features do not dictate provirus activity

To address whether integration site features affected the viral gene expression patterns observed here, zip coded provirus integration sites were compared for clones with differing expression patterns. Integration sites were determined using a linker-mediated nested PCR strategy applied to genomic DNA from the original Jurkat pool. Primers were designed so that sequencing reads included U3 resident zip codes. Initial analysis indicated variable rates of assignment of a single zip code to multiple genomic locations, likely reflecting the formation of chimeric molecules during PCR [35, 36]. We therefore went on to implement a greedy strategy to assign genomic locations to each zip code based on the number of independent DNA fragments (based on unique DNA shear points) and total reads supporting each site. Using this approach, we assigned a genomic location to each of the 225 high abundance zip codes (Supplementary Table 2). As expected [37], integrants were substantially enriched for annotated genes and genes expressed in Jurkat cells (Supplementary Figure 5), with 58% having the same orientation as the intersecting transcript (109 of 188 that intersect with single genes, p=0.034, binomial test).

To search for factors that may affect set point expression levels, we assigned each of the 225 zip codes to one of three classes: those with a GFP+ proportion of at least 0.6 in both pools (‘mostly GFP+’; 157 clones), those with a GFP+ proportion less than 0.4 in both pools (‘mostly GFP-’; 48) and those with mixed levels of GFP expression (‘mixed’; 20). Ignoring integrants that intersect with no genes or with genes having overlapping expression in divergent directions, we found no orientation preference for the ‘mostly GFP+’ integrants (65 of 129 with single intersection have same orientation; p= 0.99 binomial test), whereas both the mostly GFP- and mixed populations were enriched for integration in the same orientation as gene transcription (30 out of 40; p=0.002 and 15 out of 19; p=0.019). The GFP+ proportion of each integrant had a strong negative correlation with original abundance in the pool (Spearman ρ=-0.289, p=1.08*10^-5^).

We additionally compared the distance of integrants to enhancer associated (H3K27ac) and repressive (H3K9me3) chromatin marks previously determined in Jurkat cell lines [38, 39]. Distance to H3K27ac peaks had a negative but non-significant correlation to GFP+ proportion (Spearman ρ=-0.105, p=0.118). Distance to existing H3K9me3 repressive marks in Jurkat cells was also negatively correlated with GFP+ proportion (Spearman ρ=-0.195, p=0.0034). Thus, these results conflictingly showed that integrants with higher GFP expression states were on average closer to both existing repressive and enhancer chromatin marks. Comparing values across classes revealed the modest nature of these enrichments (Figure 7), with original clonal abundance and distance to existing H3K9me3 peaks showing a significant difference between mostly GFP+ and mostly GFP-clones (p=0.0046 and p=0.0073 Mann Whitney U 2-sided test).

**Figure 7.**
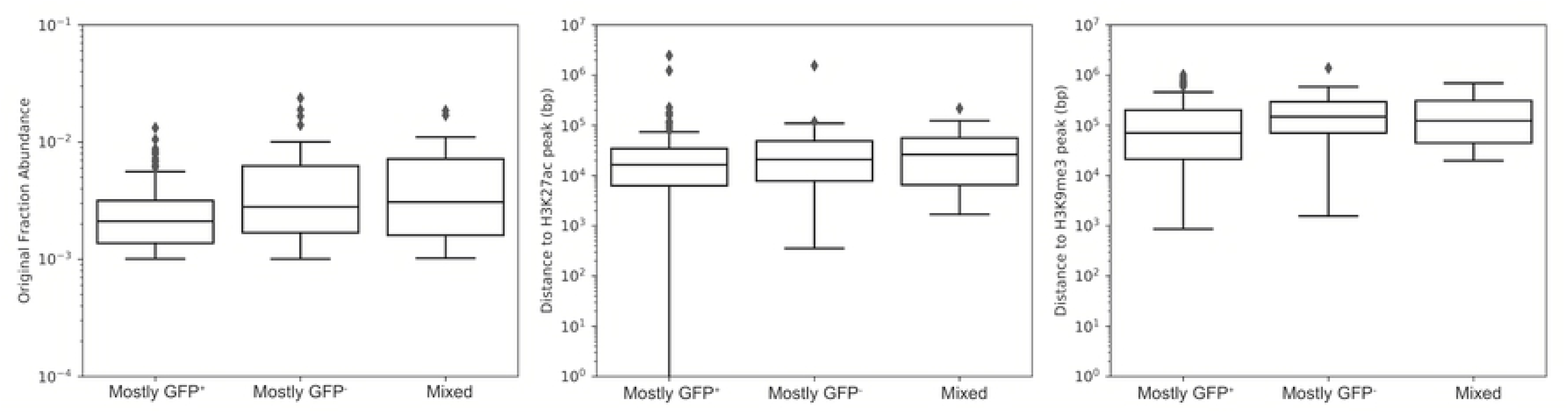
Correlations between GFP+ proportions and specific genomic features. Each of the 215 zip codes were binned into one of three categories (mostly GFP+, mostly GFP-, or mixed, as described in the text). Panel A box plots show the fractional abundance of each zip code residing in that category of clones, as determined in the original unsorted pool (Figure 2 data). Panels B and C compare distances to H3k27ac and H3k9me3 peaks, respectively, for the mostly GFP+, mostly GFP-, and mixed expression pattern zip codes. For each boxplot the median and interquartile range is depicted.

## Discussion

Here, persistence and HIV-1 expression profiles of individual integrant clones were compared within polyclonal populations using “zip coded” proviruses, each tagged to identify the genomic neighborhood where the provirus had integrated. The results revealed a complex array of heritable differences among clones in population sizes and expression characteristics.

Marking libraries with randomized sequence tags has been used in many systems including SIV and HIV-1 [40] [41] [14]. One group reported infectious SIV derivatives barcoded to track population dynamics during treatment and rebound [41]. Unlike those SIV derivatives [41], our vectors lacked Env and (except when remobilized by pseudotyping) were limited to single replication cycles. Barcodes were inserted toward the center of the virus in the SIV work, while ours were inserted near provirus edges to facilitate integration site determination. Another group described barcoded HIV-derived vectors called B-HIVE, with barcodes inserted in HIV-1’s multifunctional 5’ untranslated region. [14]. We chose to leave the 5’ leader region intact because it modulates HIV-1 expression by specifying nucleosome and transcription factor binding [9], folds into a finely-balanced equilibrium of RNA elements that regulate RNA fates [42], and is highly sensitive to mutation [43]. B-HIVE vectors encode LTR-driven GFP but no virus structural proteins. In contrast, our vectors retained *gag* and *pol*, thus allowing progeny virus production and the tracking of both virions and cellular nucleic acids. B-HIVE experiments were performed at a multiplicity of infection of 0.5 and likely included dually infected clones, while we used a much lower MOI. Additionally, we assessed expression in both unsorted cell pools and in serial sub-pools sorted for LTR reporter expression, and observed both dynamic and heritable aspects of clone-specific expression not evident in the B-HIVE work [14].

We benchmarked our system using a small (74 clones) pilot study that addressed replication fidelity. Zip code abundance varied widely in this pool, as did virus release per cell. Most zip codes lost during the second cycle of replication were significantly less abundant in the expanded cell pool derived from the first round of infection than those that persisted for the second round of replication, suggesting that population bottlenecks contributed to zip code extinction. Zip code survival rates suggested a single cycle lethal mutation rate of about 10%. This is in the range predicted by findings that roughly one in three HIV-1 genomes accumulates a reverse transcriptase-generated mutation [44]. However, the majority of patients’ persistent proviruses are defective [26, 28]. Thus, our data support the notion that defective proviruses are likely more abundant *in vivo* due to selective pressures rather than to reverse transcriptase infidelity [28, 45].

Subsequent experiments were performed in Jurkat or primary T cells, using larger zip code libraries and proviruses with GFP in the *nef* open reading frame. HIV-1 proteins including Vpr and Env, which kill or inhibit cultured cells, were absent by design [46] [47]. Within the unsorted polyclonal Jurkat pool, GFP+ cells were more numerous than GFP-cells and virion release remained robust. As previously demonstrated with similar vectors, populations were readily separable into GFP+ and GFP-pools [48–50]. GFP+ pools displayed high levels of virion release while there was a near-absence of virus from GFP-cells. All abundant zip codes were reproducibly present in both GFP+ and GFP-cell sub-pools, but to widely varying extents. Using “GFP+ proportion” to represent the fraction of each clone’s cells that sorted GFP+, most clones were either “mostly GFP-“ (with GFP+ proportions ≤0.4) or “mostly GFP+” (≥0.6).

Similar to the pilot study, the number of cells per clone in Jurkat and primary CD4+ pools spanned three orders of magnitude. Although most cells in the unsorted pool were GFP+, many clones that were mostly GFP+ were comprised of relatively few cells, and the average number of cells per mostly GFP-Jurkat clone was significantly greater than for mostly GFP+ clones. This suggests that caution is appropriate when interpreting findings based on latency models that use GFP reporters and that passage cells until GFP activity largely disappears. Specifically, our results suggest that some of the apparent increases in latency over time may reflect outgrowth by clones with low GFP+ proportions rather than proviral silencing [51].

The stability of GFP+ proportions over time was addressed by re-sorting GFP+ and GFP- sub-pools that had been passaged separately. When GFP values from the secondary sorts were weighted to reconstitute the original pool, the GFP+ proportions for each clone were remarkably similar over time. Daughter cells did not always adopt a parental phenotype, but instead “flickered” between GFP+ and GFP-. It is unclear whether the flickering observed here differs from the mosaic patterns of expression that have been described previously within individual retroviral vector cell clones. In those studies, intraclonal expression variegation was interpreted to indicate integration site-dependent differences in silencing rather than alternating waves of expression [13, 52].

Heritable high levels of variation among HIV-1 integrant clones have been reported previously; however unlike the flickering we observed, within-clone HIV-1 expression level variation has appeared relatively narrow using previous approaches [13, 52]. For example, wide inter-clone variation was reported in the B-HIVE study. However, HIV-1 expression was quantified as intracellular RNA copies per cell per barcode using an unsorted cell pool, and it was assumed that every cell within a given clone expressed LTR-driven RNA to the same extent [14]. In contrast, our results suggest that at least part of the expression differences among clones reflects that each clone consists of a phenotypic mixture of cells—some that release virus and others that do not—in heritable clone-specific proportions.

What is responsible for the clone-specific stable equilibrium mixture of GFP+ and GFP-cells within each clone? Intrinsic fluctuations in transcription factor availability and other stochastic events contribute significantly to gene expression, and can cause genetically identical cells propagated under uniform conditions to display a spectrum of phenotypes [53]. The sources, regulatory mechanisms, and implications of this genetic noise are active areas of investigation [54, 55]. Phenotypic bifurcation for HIV-1 infected cells, in which intrinsic noise in Tat expression leads to the co-existence within individual integrant clones of some cells that display high levels of expression and others that display essentially none, has previously been described [56]. Transcriptional bursting from the HIV-1 promoter is a significant source of stochastic noise [57], the bursting behavior of the export factor Rev may further exacerbate noise due to Tat [58], and the phase of the cell cycle may also exert influence [59]. These and other parameters likely contributed to the broad range of GFP+ proportion set points that differentiated clones here, even though our system was carried out in transformed cells with the intention of minimizing extrinsic variability [60].

The simplest explanation for why each clone adopted a unique GFP+ proportion set point may be that multiple inputs—some stochastic and others deterministic--combined in a clone-intrinsic manner to skew the probability that a given cell would reach the Tat threshold needed for GFP expression. The deterministic components could in concept be of either host or viral origin. However, our initial pilot experiment suggests that the principle differences were not within proviral sequences, but instead of host origin. Specifically, the amount of virus release per cell differed among zip codes when all cells with the same zip code were progeny of a single integration event, but virus release per cell was fairly uniform in a second generation when zip codes were polyclonal.

To explore host contributions due to integration site features, virus/host junctions were sequenced, integration sites determined, and the characteristics of mostly GFP+ and mostly GFP-clones compared. The results indicated that mostly GFP+ and mostly GFP-clones differed significantly in proviral orientation relative to host transcription, with mostly GFP+ clones including similar numbers of proviruses in both the same and opposite orientation as host transcription but mostly GFP-clones biased toward the same orientation. This may reflect transcriptional repression, which has been reported for HIV-1 [15, 61], although one study reported an opposite orientation bias [62]. We also assessed the correlations between repressive or activating chromatin marks previously determined in Jurkat cells [38, 39] and observed modest differences in proximity to H3K9me3 marks. However, the role of host cell features in robustly discriminating latency or viral expression remains unclear. Machine learning models can differentiate high- and low-expression genes using epigenetic and genetic features [63]. However, while that problem is comparable to distinguishing expression for integration sites, such models use many thousands of training examples and tens of epigenetic features, while our analysis was restricted to hundreds of sites and a small handful of epigenetic features previously catalogued for Jurkat cells.

Speculatively, some component of the observations here may reflect epigenetic marks introduced at the time of integration: due either to stochastic events or to differences in the intracellular environment or architecture of specific integration sites. It is generally assumed that most of the latent reservoir results from the rare infection of activated cells that transition to a memory state. However, HIV-1 can enter cells at any phase of the cell cycle. Histone biogenesis is cell cycle dependent [64] and many histone post-translational modifications are faithfully introduced onto nascent strands at the time of DNA replication. Although all epigenetic marks appear regenerated within the course of a single cell generation, some marks are copied with the replication fork while others (including H3K9me3 and H3K27me3) are deposited throughout the cell cycle [65, 66]. Because HIV can infect dividing or resting T cells, and the cell’s chromatin modification machinery displays cell cycle-dependent regulation, it is possible that integration at differing phases of the cell cycle results in distinct patterns of chromatin decoration [64, 67, 68].

It seems plausible that the HIV-1 expression variation reported here may cause some of the differences among experimental models for latency [16] and that expression flickering and differential set points of expression may be a fairly common outcome during the establishment of polyclonal HIV-1 populations. As such, these properties may contribute to defining the nascent proviral populations within infected people that are subsequently culled by immune and other selective pressures. Understanding how patterns of expression that persist compare to the palette of outcomes in the absence of selection may aid efforts to identify HIV-1’s epigenetic havens, and to the design of fruitful strategies for proviral eradication.

## Materials and Methods

### Cell line propagation

293T cells were grown from a master cell bank [69] and Jurkat (Clone E6-1) cells were obtained from ATCC. Both cell lines were maintained as lab frozen stocks and validated at the time of study by tandem repeat analysis using the Applied Biosystems AmpFLSTR™ Identifiler™ Plus PCR Amplification Kit (Thermo Fisher Scientific, Carlsbad, CA). Jurkat cells were cultured in RPMI supplemented with 10% FBS (Gemini), 100 U/mL penicillin, 100 µg/mL streptomycin, 2mM glutamine and 55µM β-mercaptoethanol at 1 x 10^6^ cell/ml, while Human Embryonic Kidney (HEK) 293 T cells were grown in DMEM supplemented with 10% FBS (Gemini) and 125 µM gentamycin. Both cell lines were maintained in a 37°C incubator containing 5% CO_2_.

### Construction of zip coded vectors

All HIV-1 vectors were templated by derivatives of the NL4-3 strain plasmid NL4-3 GPP [70] or by HIV-GPV^-^, which was derived from the GKO [25] provided by Emilie Battivelli and Eric Verdin (University of California San Francisco). HIV-GPV^-^ was constructed by replacing mKO2 in GKO with the puromycin resistance gene from NL4-3 GPP. After initial work with standard two-LTR vectors, including the pilot fidelity study described here, subsequent zip coded vector preparation used single LTR versions of these vectors. For this, both vectors were modified into single “inside out” LTR forms containing the 5’ terminal 49 bases of U3 with an engineered Cla I site plus a second unique site (either Xho I or Mlu I) in U3, and inserted into pBR322 as previously described [71]. To generate zip coded HIV-1 vector templates, the single LTR plasmid versions of NL4-3 GPP and GPV- were digested with ClaI plus Xho I or Mlu I, respectively. The resulting 11.4kb HIV vector-containing fragments free of plasmid backbone were purified from agarose using QIAquick Gel Extraction Kit (Cat No./ID: 28706 Qiagen, Germantown, MD). A 304 bp zip code-containing insert fragment pool was generated by PCR using NL4-3 GPP or GPV^-^ as template, Phusion® High-Fidelity DNA Polymerase (New England Biolabs, Inc., Ipswich, MA), and primers 5’- GACAAGATATCCTTGATCTGNNNNNNNNNNNNNNNNNNNNGCCATCGATGTGGATCTACCACACACAAGGC-3’ and 5’- CGGTGCCTGATTAATTAAACGCGTGCTCGAGACCTGGAAAAAC-3’ for GPV^-^ and 5’ GTGTGGTAGATCCACATCGATGGCNNNNNNNNNNNNNNNNNNNNCAGATCAAGGATATCTTGTCTTC-3’ and 5’- ATG CCA CGT AAG CGA AAC TCT CTG GAA GGG CTA ATT CAC TCC-3’ for NL4-3 GPP.

To generate the uncloned vector template library, the 11.4 kb fragments of GPV- or HIV- GPP were joined with their cognate 304 bp zip coded partial U3 inserts via Gibson Assembly in a molar ratio of 1:5 per reaction using HiFi DNA assembly mix (New England Biolabs) following the manufacturer’s protocol. The assembled DNA was then cleaned and concentrated using Zymo Clean and Concentrator-5 kit (SKU D4013 Zymo Research, Irvine, CA), quantified by Nanodrop (Thermo Fisher Scientific), and used directly in transfections.

### Virion production

Fresh monolayers of HEK 293T cells, approximately 70% confluent, were co-transfected with 3 µg Gibson Assembly product DNA plus 330 ng pHEF-VSV-G using polyethylenimine (Polysciences, Inc., Warrington, PA) at a ratio of 1 µg total DNA to 4 µg polyethylenimine in 800 µl of 150 mM NaCl [72]. 24 hours post-transfection, DMEM was replaced with 4 ml RPMI1640 medium with 10% FBS and 1% Pen/strep. Culture supernatant was harvested at 48 hours post-transfection and filtered through a 0.22 µm filter (Fisher Scientific. Cat. No. 09-720-511). Released virus was quantified using a real-time reverse-transcription PCR assay and normalized for p24 level based on p24 protein values determined in parallel for reference samples [71]. Zip coded virus stocks were titered by infecting 90% confluent HEK 293 T cells and selecting in puromycin. Colony forming units per milliliter of viral media as determined on 293T cells was the standard for defining infectious titer in this work.

### Infection of HEK 293 T and Jurkat cells

The media on 10 cm plates of 90% confluent HEK 293 T cells was replaced with 2000 µl infection mix comprised of the indicated amount of virus-containing medium plus additional DMEM in 1 mg/ml polybrene, then incubated at 37 °C with 5% CO_2_ for 5 hours. After incubation, the infection mix was replaced with 10 ml of fresh media. Twenty-four hours post-infection, cells were placed in media containing puromycin at a concentration of 1µg/ml, which was replaced every three days for 2 weeks. Following this, colonies were individually cloned, pooled together for subsequent experiments, or stained with crystal violet and counted.

For Jurkat cell infections, virus-containing media and polybrene at a final concentration of 0.5 mg/ml were brought to a total volume of 1000 µl. This infection mixture was added to 1.5 x 10^6^ Jurkat cells and incubated in one well of a 12 well tissue culture plate (Fisher Scientific, Cat. 150628) at 37 °C with 5% CO_2_ for 5 hours. Infected cells were then transferred to Eppendorf tubes and centrifuged for 5 minutes at 2500 rpm at 4°C. Following centrifugation, supernatants were replaced with fresh media and cell pellets were resuspended and cultured at 37 °C with 5% CO_2_. At 24 hours post-infection, puromycin was added to a final concentration of 0.5 µg/ml. The infected cells were expanded into 6 cm culture plates on day 5. Ten days post-infection, the culture supernatant was replaced with fresh media and the cultures were divided into aliquots, to be either frozen or further expanded for subsequent experiments.

### Primary T cell isolation and infection

Peripheral blood mononuclear cells (PBMCs) were isolated from fresh human blood from healthy donors provided by the Department of Pathology at the University of Michigan using Ficoll Histopaque as described earlier [73]. All use of human samples was approved by the Institutional Review Board at the University of Michigan. Total CD4+ T cells were then purified from PBMCs using MACS beads (Miltenyi Biotec Bergisch Gladbach, Germany) as per the manufacturer’s instructions. On day 0, a total of 5 x 10^6^ cells were seeded in complete culture medium composed of RPMI supplemented with 10% FBS, 100 U/mL penicillin, 100 µg/mL streptomycin, 2mM glutamine and 55µM β-mercaptoethanol at 1 x 10^6^ cell/ml. The cells were stimulated using plate-bound anti-CD3 (5 µg/mL; eBioscience, Thermo Fisher Scientific) and soluble anti-CD28 (1 µg/mL; eBioscience, Thermo Fisher Scientific) antibodies in the presence of 50 U/ml IL-2 (PeproTech, Inc., Rocky Hill, NJ). On day 2 of activation, the cells were infected by spinoculation at 2500 rpm for 90 minutes at 37°C with 125 µL zip coded viral media and 0.4 mg/ml polybrene (Sigma Aldrich, St. Louis, MO) in 2.5 ml of supplemented RPMI. After spinoculation, media containing virus was replaced with fresh supplemented RPMI and cells were cultured further and expanded as needed. On day 7 post-activation, cells were harvested and sorted into GFP^+^ and GFP^-^ sub-pools by flow cytometry using FACS Aria II (BD Biosciences, Franklin Lakes, NJ) or iCyt Synergy SY3200 (Sony Biotechnology, San Jose, CA) cell sorter. A portion (2 x 10^6^ cells) of GFP^-^ CD4+ T cells were subjected to puromycin selection for another 48 hours using 1ml supplemented RPMI containing IL-2 and 2 µg/ml puromycin and then harvested for further analysis.

### Flow cytometry

For flow cytometry analysis and sorting, Jurkat or primary T cells were suspended in phosphate buffered saline (PBS) containing 1% FBS (FACS buffer). Dead cells were excluded in all analyses and sorting experiments using propidium iodide (PI). Intracellular Gag staining was carried out using a Gag monoclonal antibody conjugated to Phycoerythrin (KC-57 RD1 Beckman Coulter). 1×10^5^ cells from a HIV GPV-zip coded library were washed once with FACS buffer and fixed with 100 µl of BD cytofix for 10 minutes at room temperature in the dark. Cells were then washed twice with FACS buffer then once with BD perm/wash buffer. Staining was carried out at a 1:200 dilution of antibody in 1x BD perm buffer. The cells were incubated in the dark at room temperature for 15 minutes, washed twice, then resuspended in 200 µl FACS buffer. Acquisition was carried out on the FITC channel for GFP and PE channel for Gag. Cell fluorescence was assessed using FACSCanto II (BD Biosciences) and data were analyzed using FlowJo software, version 9.9 (FlowJo, LLC., Ashland, Oregon).

### PCR amplification of zip codes from zip coded cells and virus

Genomic DNA was extracted from zip coded cell libraries using Qiagen DNeasy Blood & Tissue Kit (Qiagen, Germantown, MD). Zip codes were amplified from 100 ng of genomic DNA using primers flanking the zip code region (primers: 5‘-NNACGAAGACAAGATATCCTTGATC-3’ and 5’-NNTGTGTGGTAGATCCACATCG-3’) using Phusion® High-Fidelity DNA Polymerase (New England Biolabs) in HF Buffer. For zip code amplification, we designed multiple primers complementary to the template binding site that included two known, random nucleotides at the 5’ end for use in separate reactions. By comparing the primers used for amplification and the nucleotides at the end of each amplicon, we could confirm that PCR cross contamination had not occurred. Reactions were cycled 26-35 times with 30 second extension at 72^0^ and a 59^0^ annealing temperature. Zip coded amplicons were purified with DNA Clean & Concentrator-5 (Zymo Research, CA. Cat. No. D4013) and eluted in 20µl of H_2_O. To amplify zip codes from virus, virus-containing media was filtered through a 0.22 µm filter, concentrated by ultracentrifugation at 25,000 rpm through a 20% sucrose cushion, and RNA extracted with Invitrogen TRIzol Reagent (Thermo Fisher Scientific). The dissolved RNA was treated with RQ1 DNase (Promega, Fitchburg, WI) to remove possible DNA traces, re-extracted with phenol-chloroform, and stored at −80 °C. cDNA was synthesized using M-MLV RT (H–) (Promega) and U3 antisense primer 5’-TGTGTGGTAGATCCACATCG-3’. Zip codes were amplified from this cDNA using conditions outlined above.

For library construction, protocols and reagents from NEBNext® Ultra™ DNA Library Prep Kit for Illumina® (New England Biolabs) were used for end repair, dA-tailing, and to ligate Nextflex adapters (Perkin Elmer, Waltham, MA) onto amplicons. After ligation, reactions were diluted up to 100 µl with H2O, purified with 0.85x SPRIselect beads, washed twice in 70% ethanol, and eluted into H_2_O. PCR enrichment of adapter-ligated amplicons was done for 7 cycles using NEBNext® Ultra™ DNA Library Prep Kit, reactions were diluted up to 100 µl with H_2_O, and purified with 0.85x SPRIselect beads (Beckman Coulter) as outlined above. Libraries were quantitated with KAPA Library Quantification Kits for Next-Generation Sequencing (Roche Sequencing Solutions, Inc., Pleasanton, CA) and Qubit™ dsDNA HS Assay Kit (Thermo Fisher Scientific), pooled equally, and sequenced with MiSeq Reagent Kit v3, 150 cycle PE on MiSeq sequencer (Illumina, San Diego, CA).

### Calculating GFP+ proportions

GFP+ proportions were calculated by dividing the read frequency of each zip code within GFP+ sorted cells by the zip code’s summed abundance in both GFP+ and GFP-sorted cells, after weighting values to reflect the GFP+ and GFP- sub-pools’ fractions of total cells. For example, a clone’s GFP+ read frequency would be the proportion of GFP+ total reads that contained that clone’s zip code. If the total pool was 75% GFP+ and 25% GFP-cells, a clone’s weighted abundance would be three times its abundance in GFP+ cells plus its abundance in GFP-cells. Thus, if a given zip code were 2% of the GFP+ cells and 3% of the GFP-cells within a 75% GFP+/25% GFP-total pool, its GFP+ proportion would be 2% divided by [(3 x 2%) + 3%] or 22%.

### HIV integration-site sequencing

Template for hemi-specific ligation mediated PCR of insertion sites was obtained by linear PCR and biotin enrichment of sheared, genomic DNA with linkers ligated on each end. Linker was synthesized by mixing oligo 5’– GTAATACGACTCACTATAGGGCTCCGCTTAAGGGACT-3’ and 5’–PO4-GTCCCTTAAGCGGAG-3’-C6 [74] at a final concentration of 40 µM each in 100 µl volume. Oligo mixture was heated in PCR block for 5 minutes at 95°C, PCR machine was immediately shut off, and block was allowed to cool for 2 hours to room temperature. Genomic DNA was extracted from cells using Qiagen DNeasy Blood & Tissue kit (Qiagen) and 200 ng of DNA was sheared to 1 kb fragments using Covaris M220 and micro-TUBE according to manufacturer’s recommended settings (Covaris, Woburn, MA). Sheared DNA was purified with 1x SPRIselect beads according to manufacturer’s instructions (Beckman Coulter) and sheared ends were repaired with NEBNext® Ultra™ End Repair/dA-Tailing Module (New England Biolabs) according to manufacturer’s protocol. Repaired, dA-tailed DNA was purified with 0.7x SPRIselect beads (Beckman Coulter) and the partially double stranded DNA linker with dT overhang was ligated in a 60 µl reaction containing 6ul of 10X T4 DNA Ligase Buffer, 1.33 µM linker DNA, and 3600U Ultrapure T4 DNA ligase (Qiagen) at 16°C for 16 hours followed by 70°C incubation for 10 minutes. Ligated DNA was purified with 0.7 x SPRIselect beads (Beckman Coulter) and used for template in linear PCR reaction containing 1x Expand Long Range Buffer, 500 µM dNTPs, 3% DMSO, 3.5U Long Range Enzyme Mix, and a 500 µM biotinylated primer that anneals to the HIV LTR in our construct, 5’- /52-Bio/CAAAGGTCAGTGGATATCTGACCCC-3’. Cycling parameters were 95^0^ C for 5 minutes, 40 cycles of 95^0^ C for 45 seconds, 60^0^ C for 1 minute, and 68^0^ C for 1.5 minutes, followed by a 10 minutes incubation at 68^0^ C. PCR product was purified with 1x SPRIselect beads (Beckman Coulter), resuspended in 20 µl H2O, and biotinylated fragments were captured using Dynabeads kilobase BINDER kit (Thermo Fisher Scientific) according to manufacturer’s instructions. DNA captured by beads was used as template in a hemi-specific PCR reaction containing 1x Expand Long Range Buffer, 500 µM dNTPs, 3% DMSO, 3.5U Long Range Enzyme Mix, 500 µM of a nested primer that anneals to HIV LTR in our construct, 5’- GCCAATCAGGGAAGTAGCCTTGTGTGTGG-3’, and 500 µM of a primer that anneals to the linker, 5’-AGGGCTCCGCTTAAGGGAC-3’. Cycling parameters were 95^0^ C for 5 minutes, 30 cycles of 95^0^ C for 45 seconds, 60^0^ C for 1 minute, and 68^0^ C for 1.5 minutes, followed by 10 minutes’ incubation at 68^0^ C. PCR product was purified with 0.7x SPRIselect beads (Beckman Coulter), then protocol and reagents from NEBNext® Ultra™ DNA Library Prep Kit for Illumina (New England Biolabs) were used to end repair, dA-tail, and ligate Nextflex sequencing adapters (Perkin Elmer) onto amplicons. Ligation reaction was purified with 0.65x SPRIselect beads (Beckman Coulter) and 7 cycles of PCR to enrich for ligated product was done with NEBNext® Ultra™ DNA Library Prep Kit for Illumina (New England Biolabs). Libraries were quantitated with KAPA Library Quantification Kits for Next-Generation Sequencing (Roche Sequencing Solutions, Inc., Pleasanton, CA) and Qubit™ dsDNA HS Assay Kit (Thermo Fisher Scientific), pooled equally, and sequenced with MiSeq Reagent Kit v3, 600 cycle PE on MiSeq sequencer (Illumina, San Diego, CA). All generated sequence data has been deposited to the Sequence Read Archive (SRA) under project accession PRJNA531502

### Zip code analysis and quantification

Zip codes were identified and quantified from Illumina sequencing reads using a custom suite of tools implemented in Python (https://github.com/KiddLab/hiv-zipcode-tools). First, 2×75 bp paired reads were merged together using *flash* v1.2.11 [75]. Zip codes were identified by searching for known flanking sequence (with up to 1 mismatch). Only candidate zip codes with a length of 17-23 nucleotides were considered and the number of read count for each unique zip code was tabulated. To identify the set of zip codes for further analysis, zip codes families which account for PCR and sequencing errors were determined by clustering together the observed unique zip codes. Comparisons among zip codes were calculated using a full Needleman-Wunch alignment tabulated with a score of +1 for sequence matches, −1 for mismatches, and a constant gap score of −1. Comparisons with two of fewer mismatches (counting a gap as a mismatch) were accepted as a match. Using this criteria clusters were then identified. First, unique zip codes were sorted by abundance. Then, beginning with the most abundant zip code, each sequence was compared with all of the previous zip codes. If no previous zip code had two or fewer mismatches that zip code was accepted as a cluster and then the next most abundant zip code was considered. This process was continued until the first unique zip code having a match to a more abundant zip code was identified. This defined the set of families for consideration. Abundance for the families was then determined by assigning unique zip codes to the most abundant family whose sequence was within 2 mismatches and summing their associated read counts.

In sorting experiments, the GFP+ proportion for each zip code was determined as F_i_ = (G_i_ * P)/ (G_i_ * P + W_i_ * Q) where F_i_ is the GFP+ fraction of zip code i, G_i_ is the fraction abundance of zip code i in the GFP+ sorted pool, W_i_ is the fraction abundance of zip code i in the GFP-sorted pool, P is the fraction of cells that sorted into the GFP+ pool and Q is the fraction of cells that sorted into the GFP-pool. In the Jurkat pool 1, the initial GFP+ fraction was 0.524 and the initial GFP-fraction was 0.36. Of the GFP+ sort from pool 1 the GFP+ fraction was 0.887 and the GFP-fraction was 0.079 GFP-while in the GFP-sort from pool 1 the GFP+ fraction was 0.046 and the GFP-fraction was 0.928. In the Jurkat pool 2, the initial GFP+ fraction was 0.518 and the initial GFP-fraction was 0.364. Of the GFP+ sort from pool 2 the GFP+ fraction was 0.915 and the GFP-fraction was 0.082 GFP-while in the GFP-sort from pool 2 the GFP+ fraction was 0.063 and the GFP-fraction was 0.923. For primary cell data analysis, the abundance of each zip codes in the GFP+ and GFP-pools summed, and only those zip codes with summed abundance greater than 0.001 in both replicates were considered, and a GFP+ fraction of 0.9 and a GFP-fraction of 0.1 were assumed.

Analysis of integration sites occurred in two stages. First, read-pairs were analyzed to identify which read derived from the LTR sequence and which from the genomic linker. Zip code sequences were extracted from the LTR-derived read based on matches to flanking sequence in the vector as described above. The linker sequence and LTR sequence flanking the zip code were removed and the extracted zip code sequence was then associated with the remaining portion of each read pair. Second, the trimmed read pairs were aligned to a version of the hg19 genome that included the sequence of the utilized HIV vector using bwa mem version 0.7.15. The resulting alignments were then parsed to identify the shear point (DNA adjacent to where the linker was ligated) and integration point (the DNA location adjacent to the LTR sequence). The zip codes were then assigned to previously identified zip code families, and the number of unique shear points and total reads supporting a integration site for each zip code were tabulated. Only reads with a mapping quality greater than 10 were considered. A greed algorithm was then used to associate each zip code with a genomic location. Candidates assignments were sorted by the number of shear points and total reads in descending order. The most abundant assignment was taken as the position for the indicated zip code, other assignments for that zip code were removed, and the process was repeated.

### Determination of chromatin marks and expressed genes

Gene annotations were determined based on Ensembl release 75. Jurkat gene expression data produced by Encode [76] was used (accession ENCSR000BXX), and genes with TPM counts greater than 5 in both replicated were considered to be expressed. H3K27ac peaks were identified using data from [38] (GSM1697880 and GSM1697882). Chip-seq and control data were aligned to hg19 using bwa mem and peaks were identified using macs2 v [77] with the --nomodel option. For H3K9me3 peaks, data from [39] (GSM1603227) were aligned to hg19 using bwa mem and processed using macs2 without a control sequence set. For both marks a p value cutoff of 1*10^-9^ was used.

### Ethics statement

Peripheral blood mononuclear cells (PBMCs) were isolated from fresh human blood from healthy donors provided by the Department of Pathology at the University of Michigan. All samples were anonymized and all use of human samples was approved by the Institutional Review Board at the University of Michigan

## Figure Legends

Figure S1. **Zip code family and read abundance for single cycle experiment.** The blue line (left axis) shows the number of unique zip code families determined by clustering the indicated number of unique zip codes. The red line (right axis) show the cumulative fraction of reads accounted for by each unique zip code.

Figure S2. **Zip code rank and fractional abundance for Jurkat pool.** Axes are as in Figure S1.

Figure S3. **Flow cytometric analysis for the co-occurrence of intracellular Gag staining and GFP.** Performed using Jurkat cells containing zip coded HIV GPV-library as described in Materials and Methods. Numbers in each quadrant indicate the proportion of total cells in that quadrant.

Figure S4. **Observed abundances for zip codes from primary cells infections.** The Y axis shows the mean fractional abundance for each zip code across the two parallel sub-pools.

Figure S5. **Enrichment of integrants in annotated genes (left) and genes expressed in Jurkat cells (right).** Observed intersections compared with random placements. In each figure the red line indicates the observed values for the 225 integrants determined in the Jurkat cells and the blue histogram indicates the counts observed from 1,000 random permutations.

